# The effects of LED daylength extensions on the fecundity of the pest aphid *Myzus persicae* and the daily activity patterns of its parasitoid, *Aphidius matricariae*

**DOI:** 10.1101/2023.03.12.532303

**Authors:** Jessica L. Fraser, Paul K. Abram, Martine Dorais

## Abstract

Supplemental lighting, such as with LED lamps, allows greenhouse producers to maintain yields when natural light levels are low. Since both insect pests and their natural enemies are sensitive to light, both the duration and spectrum of LED daylength extensions could affect biological pest control in greenhouses. Longer days could allow for extended periods of reproduction for pests or foraging activity of biological control agents, possibly depending on the spectra used for these extensions. However, the effects of lengthening days with different LED spectra on the behaviour of biological control agents has mostly been studied in short-term experiments to date, and has not always included the context of the light’s effect on their hosts’ reproduction. In growth chambers, we examined the locomotor activity of the parasitoid biocontrol agent *Aphidius matricariae* (Hymenoptera : Braconidae) as a predictor for foraging activity and the fecundity of its aphid host *Myzus persicae* (Hemiptera : Aphididae) over multiple days under different daylength extension regimes representative of those used in greenhouse vegetable production. We compared the effects of 14, 16, 18, and 20 h photoperiods, and 12 h days extended by 6 h with three different spectral qualities. The parasitoids adjusted how their activity was distributed throughout the lit period of the day (i.e., its total duration and peak timing) without changing the total amount of daily activity, regardless of the photoperiod or the light spectrum used for daylength extension. The aphids’ peak fecundity was not affected by photoperiod or spectral quality. Our results suggest that at least some behavioral and reproductive traits of these insects can be resilient to even drastic changes in their light environment.

## 1. Introduction

Supplemental lighting is beneficial in vegetable greenhouses for maintaining yields when natural light levels are low, allowing producers to maintain adequate daily light integrals even in winter in northern climates (Dorais, 2003; Dorais et al., 2017). While HPS lamps have long been the most popular choice, LEDs are promising because they are energy-efficient, they can be placed closer to plants, and their spectral quality is customizable (Hao et al., 2018; Singh et al., 2015). The spectral quality used influences the growth and nutritional quality of the crops, as well as allowing producers to maximize energy efficiency by selectively producing the most optimal wavelengths (Olle and Viršile, 2013).

While LED usage is typically tailored to the needs of greenhouse plants, there are often other organisms present in greenhouses that have the potential to be impacted by lighting conditions. Arthropod biocontrol agents are widely employed for pest management in greenhouses (van Lenteren et al., 2018), and both they and their target pests can respond to changes in the greenhouse light environment (Johansen et al., 2011). Diurnal natural enemies could benefit from lengthened photoperiods that allow them more time to forage for hosts or prey (Kehoe et al., 2020). Indeed, longer days do produce greater parasitism or predation rates in some diurnal or crepuscular species (Joschinski et al., 2019; Kehoe et al., 2020; Kehoe and van Veen, 2022; Zilahi-Balogh et al., 2006) with longer nights similarly able to benefit nocturnal species (Joschinski et al., 2019). The color of light used to extend the day is also relevant to their foraging, since spectral quality can influence natural enemies’ attraction into a desired area (Park and Lee, 2021), locomotor activity (Cochard et al., 2017; Wang et al., 2013), host encounter probability (Cochard et al., 2019a), and parasitoid sex ratios (Cochard et al., 2019b). Under the right circumstances, it is possible to manipulate lighting conditions in ways that do not affect pests and biocontrol agents equally. For example, excluding UV from a greenhouse can impede aphid and whitefly dispersal without reducing the host-finding abilities of some aphid parasitoids (Chiel et al., 2006; Chyzik et al., 2003; Dáder et al., 2017). It may therefore be possible to selectively benefit natural enemies through the choice of greenhouse lighting.

Despite its potential impacts, the effects of LED greenhouse lighting—especially in terms of spectral quality—on parasitoid biocontrol agents have been studied very little to date (Cochard et al., 2019b; Johansen et al., 2011; Lazzarin et al., 2020), and the existing research on how LED spectra affect parasitoid behavior have based their conclusions on very short-term observations (Cochard et al., 2017; Cochard et al., 2019a; Cochard et al., 2019b). Given that greenhouse biocontrol agents typically live for multiple days, it is important for them to be able to adapt to the lighting conditions that they encounter. However, to our knowledge, it is not known how parasitoids adapt their daily foraging patterns in response to greenhouse light conditions incorporating novel spectra, and whether their sensitivity to novel spectra remains consistent or diminishes over time. We sought to examine the effects of the photoperiod and spectral quality of LED lighting on a parasitoid biocontrol agent’s foraging over a longer period, while also accounting for effects the light might have on reproductive output of their aphid hosts. This information could provide insights into whether crop lighting regimes could be tailored to manipulate parasitoid behavior and improve greenhouse biocontrol.

In this study, we examined in growth chambers the effects of LED-extended days on a parasitoid’s locomotor activity and its aphid host’s fecundity under a range of artificially-lengthened photoperiods and days lengthened with different spectra. For our pest insect, we used the aphid *Myzus persicae* (Hemiptera: Aphididae), which is a serious greenhouse pest (Blümel, 2004). Its visual system has been well characterized, and it has photoreceptors with peak sensitivities at about 330 to 340 nm (UV-A), 490 nm, (blue), and 530 nm (green) (Kirchner et al., 2005). For our natural enemy, we studied the parasitoid *Aphidius matricariae* (Hymenoptera: Braconidae), a commercially-available biocontrol agent used to manage *M. persicae* in pepper greenhouses (Blümel, 2004). This parasitoid presumably also has photoreceptors with peak sensitivity to UV, blue, and green, which is typical in wasps and bees (Peitsch et al., 1992).

We examined aphid reproduction in terms of the number of offspring produced during the peak reproductive period as a predictor for aphid population growth once the aphid has settled on the plant. For the parasitoid, we examined spontaneous locomotor activity as a proxy for foraging activity and ultimately parasitism, since locomotor activity timing has previously been shown to be correlated with parasitoid oviposition patterns (Fleury et al., 2000). We hypothesized that parasitoids would adjust their activity to remain active under longer days and show increased foraging activity overall. We also hypothesized that they would be more active under broader spectra containing greater proportions of the short- to mid-length (UV, blue, and green) wavelengths they are sensitive to. We hypothesized the aphids would show at most a modest increase in fecundity in response to lengthened days or more long-wavelength-enriched spectral qualities, as these conditions could influence their circadian rhythms (Joschinski et al., 2015) or favor feeding behaviour (Fennell et al., 2020), though not necessarily enough to provoke significant changes in population size. Thus, we predicted that it may be possible to find a combination of photoperiod and spectral quality that produces a greater benefit to the parasitoids’ total foraging activity than to their hosts’ reproduction.

## 2. Materials and methods

### 2.1 Plant production

Plant material used in rearing and experiments was grown from seed in greenhouses at the Agriculture and Agri-Food Canada Agassiz Research and Development Centre (AAFC ARDC; 49°14’33.0” N, 121°45’45.2” W). *Capsicum annuum* var. Red Bell Pepper (McKenzie seeds, Brandon, Canada) was used for the photoperiod experiments, and var. Seminis hybrid sweet pepper PS 0941819 (Bayer, Leverkusen, Germany) for the spectral quality experiments. Plants were grown in a peat and perlite mix medium (pH 5.6–6.2) and hand-watered daily with approximately 100 mL of an 18N-6P-20K complete solution (Poinsettia Plus, Master Plant-Prod, Brampton, Canada) (pH 6.0–6.4) containing 100 ppm total N, 14 ppm P, 93 ppm K, 11 ppm Mg, 0.560 ppm Fe, 0.280 ppm each of Zn and Cu, 0.0560 ppm B, and 0.336 ppm Mo. The plants were provided fresh water once per week to prevent fertilizer buildup, and were covered by mesh insect cages to protect them from pests. They were grown under natural light supplemented with 400–600W HPS lamps (Model LR48877, P.L. Light Systems, Beamsville, Canada; Model LR86176, Light-Tech Systems, Stoney Creek, Canada). The lamps provided approximately 50 μmol m^-2^ s^-1^ photosynthetic photon flux density (PPFD) (average 47.9 μmol m^-2^ s^-1^, standard error [SE] 7.2, from 10 readings) at canopy height, after accounting for absorption by the cages. Light readings were taken using an Optimum SRI-2000 handheld spectrophotometer (Optimum OptoElectronics Corp, Hsinchu, Taiwan). The lamps were on for a 16 h photoperiod, and turned off when sun intensity was at or above 400 W m^-2^. Greenhouse target temperatures were 20–22°C during the day and 18–20°C at night, with relative humidity maintained between 40 and 65%. Plants were used in experiments at 58–59 days old. This age was chosen to provide sturdy leaves that would remain in good condition for long periods after being removed from the plant, even with frequent handling.

### 2.2 Insect rearing

#### Aphids

We used *M. persicae* from a single-clone line maintained at the AAFC ARDC (Uriel et al., 2021). This aphid has a green and a pink morph, but we exclusively used green apterous individuals in experiments. Colonies of *M. persicae* were maintained indoors in dedicated rearing rooms under fluorescent lights (Model F32T8, Philips Lighting, Eindhoven, Netherlands) providing a mean of 40.2 μmol m^-2^ s^-1^ (SE 2.45) PPFD at canopy height (from 10 readings with an Optimum SRI-2000 handheld spectrophotometer) with a 16L:8D photoperiod on whole pepper plants at 20 °C and at 40–60% relative humidity. Aphids used in experiments were specially reared to ensure they were the same age and experienced similar levels of crowding. Individual aphids from the main colony were removed and placed into individual 120 mL polystyrene coffee cups (Loblaw Co. Ltd, Brampton, Canada) which contained a pepper leaf trimmed to fit inside. At the bottom of each cup was approximately 45 mL of water covered by a 59 mL plastic SOLO cup (Dart Container Corporation, Toronto, Canada), and the cups were covered with tissue affixed with a plastic coffee cup lid (Dixie, Atlanta, USA). One aphid was placed into each cup with a paintbrush and allowed to reproduce. The oldest offspring were collected 6 days later to use in experiments.

#### Parasitoids

We founded *A. matricariae* colonies from a starting supply of about 500 parasitoids that were purchased from Biobest (Biobest Group NV, Westerlo, Belgium), and replenished it in the same way every four to five months to avoid potential inbreeding depression. The parasitoids were maintained on *M. persicae* from the main aphid colony under fluorescent light (Model F32T8, Philips Lighting, Eindhoven, Netherlands) providing a mean of 27.4 μmol m^-2^ s^-1^ (SE 4.45) PPFD at canopy height (based on 10 readings with an Optimum SRI-2000 handheld spectrophotometer) with a 16L:8D photoperiod at 20°C and 40–60% relative humidity. For each generation of parasitoids, multiple cages were prepared as follows: 24 adult aphids were divided over four plants, and after five days—to allow the aphids to produce nymphs—20 adult parasitoids were released into the cage. The parasitoids were provided with drops of honey for feeding. When mummies developed (9–11 days after parasitoids were introduced), they were carefully removed from the plant and transferred into a new cage containing a pepper plant, honey droplets, and a capped, water-filled cup with a cotton wick. Age-synchronized parasitoids for experiments were collected on the day of their emergence by capturing them carefully in a clean glass pipette. Parasitoids were sexed by visual inspection of the abdomen (Giri et al., 1982). Sexed parasitoids (< 24 h old) were placed into transparent ventilated plastic containers (960 mL) (BugDorm insect pot, MegaView Science Co., Ltd., Taiwan) containing a pepper leaf and a cotton wick in a cup of water and drops of honey. Six females and six males were put in each cup, and left until the following day to allow them to mate.

Voucher specimens for both species of insects used are preserved at the AAFC ARDC, Agassiz, BC, Canada.

### 2.3 Growth chamber experiments

In order to study the effects of photoperiod and of LED day extensions of varying spectral qualities on the aphid and parasitoid, we conducted two growth chamber experiments from April 15th to July 19th 2021, with trials running simultaneously in the same growth chambers so that they experienced identical conditions. We performed two temporal replicates for both the photoperiod and spectral quality experiments, randomly assigning light treatments among the growth chambers in each replicate.

Each of four growth chambers (Conviron Adaptis, Controlled Environments Limited, Winnipeg, Canada) was equipped with two programmable Heliospectra RX30 programmable LED grow lights (Heliospectra Canada Inc., Toronto, Canada) suspended at 78.1 cm above a rack upon which specimens were placed. Beneath the lamps, 10 layers of white polyester fabric (Reemay, Avintiv, USA) were suspended across the growth chamber to dim the light. Different light settings were used depending on the experiment (Figure 1). The light intensity at approximately insect height was, on average, 87.4 μmol m^-2^ s^-1^ total photon flux density (250–800 nm) for the broad-spectrum white (BSW) treatment, 89.2 μmol m^-2^ s^-1^ for the red and blue (RB) treatment, and 83.3 μmol m^-2^ s^-1^ for the red, green, and blue (RGB) treatment as measured with an Optimum SRI 2000 handheld spectrophotometer. Photon flux distribution plots (Figure 1) were prepared using the R package *pavo* (Maia et al., 2019). Further details on the light treatments, including the proportion of each wavelength range, are provided in the Appendix (Table A.1). The temperature was set to maintain a constant 20°C (mean ± SE for each growth chamber: 20.1°C ± 0.001, 20.2 °C ± 0.002, 20.1°C ± 0.002, 20.3°C ± 0.002). Relative humidity was maintained between 60 and 85% (mean ± SE for each growth chamber : 79.3% ± 0.02, 69.3% ± 0.02, 73.9% ± 0.02, 68.5% ± 0.02), from data recorded using HOBO U23 Pro v2 loggers, Onset Computer Corporation, Bourne, United States). As a consequence of sharing a growth chamber, insects within the same light treatment in a given temporal replicate were not completely independent (Rogers et al., 2021), though we tried to mitigate this by making the growth chambers as uniform as possible and randomly assigning light treatments to different growth chambers in each temporal replicate.

**Figure 1.**
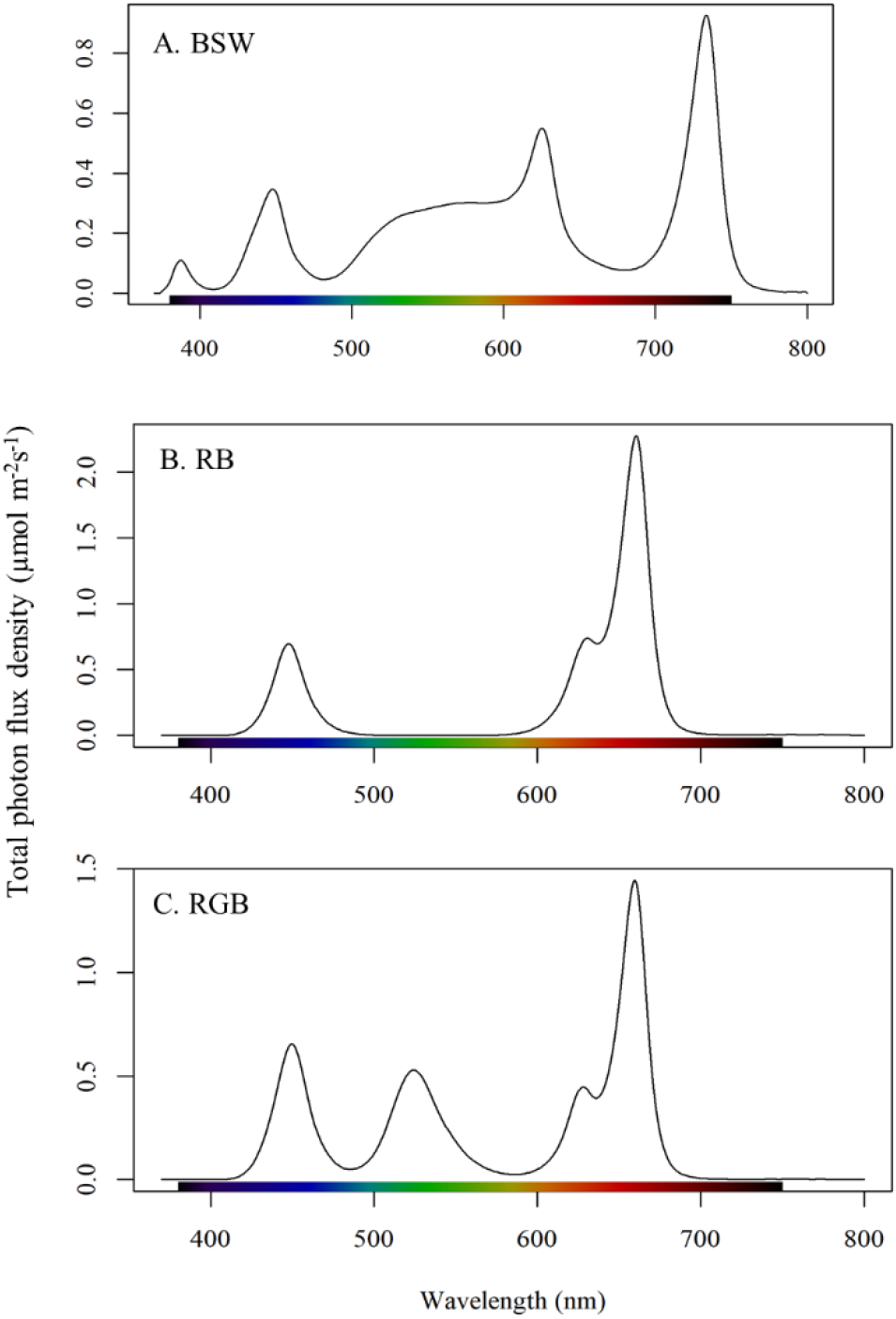
Spectra used in photoperiod and spectral quality experiments. (A) The BSW treatment (“BSW”) was composed of a 5700K white LED plus narrowband LEDs with peaks at 380 nm (UV), 620 nm, and 735 nm (far red). (B) The red and blue treatment (“RB”) was composed of narrowband LEDs with peaks at 450 nm (blue), 620 and 660 nm (red). (C) The red, green, and blue treatment (“RGB”) was composed of LEDs with peaks at 450 nm, 530 nm (green), 620 and 660 nm.

### 2.4 Experiment 1: The effect of photoperiod on aphid reproduction and parasitoid activity

In the photoperiod experiments, we sought to examine how the duration of supplemental lighting used in greenhouse production affects aphid reproduction and parasitoid locomotor activity. For all four photoperiod treatments, lights turned on at 04:00 with the BSW setting (which represents a 2–hour phase shift relative to the insect rearing rooms). They turned off at 18:00, 20:00, 22:00 or 23:59, producing 14L:10D, 16L:8D, 18L:6D, and 20L:4D photoperiods respectively. These photoperiods were selected to cover, in equal increments, the range that might be used in a commercial greenhouse. It is most common for producers to use a 14–17 hour photoperiod for peppers and other fruit vegetable crops (Hao et al., 2018), but healthy peppers can be grown under photoperiods as long as 20 hours (Demers et al., 1998), depending on the light conditions. The same light setting was used for all treatments: a broad-spectrum white (hereafter, BSW), designed to include wavelengths that are present in sunlight, including UV-A and far-red (Figure 1). The BSW treatment consisted of a 5700K white LED supplemented with narrowband LEDs with peaks at 380 nm (UV-A), 620 nm (red), and 735 nm (far-red). The mean intensity of the BSW light was 86.5–89.8 μmol photons m^-2^ s^-1^ including all wavelengths from 250–850 nm.

#### Aphid reproduction

In order to study whether aphid reproductive output varies with photoperiod, an experiment was conducted with aphids in separate containers on individual pepper leaves. To prepare the experimental containers (Figure A.1), we removed large mature leaves from 58 or 59-day-old plants at the base of the petiole, and inserted the petiole into a covered cup of water through a hole in the lid. The hole was further plugged with a piece of a cotton dental roll. Then the whole was put into a transparent plastic cup (960 mL) with a mesh-covered lid (BugDorm insect pot, MegaView Science Co., Ltd., Taiwan). The pepper leaf was gently folded down to fit into each container, such that the adaxial side faced the lid. For each leaf, three individual 6-day-old aphids were gently placed onto the leaf with a paintbrush. The leaves, rather than the aphids, were considered the experimental units. Forty of these containers were prepared for each temporal replicate, with 10 randomly assigned to each of four growth chambers. In total, 80 leaves were used over the two temporal replicates of each experiment, with 20 in each light treatment. Aphids were counted every Monday, Wednesday, and Friday. For counting, cups were removed from the growth chambers. Each leaf was carefully unfolded and all aphid nymphs in the container were counted and removed with a moistened paintbrush. Each replicate of the experiment was run for 18–19 days, to capture the aphids’ peak reproductive period (Uriel, 2020, Supplementary material).

#### Parasitoid locomotor activity and lifespan

To determine how parasitoid locomotor activity levels, as a predictor of foraging activity and ultimately parasitism rates, vary with photoperiod, experiments were designed and conducted using TriKinetics LAM25H infrared activity monitors (TriKinetics Inc., Waltham, USA). This method has been used previously to assess locomotor activity as a proxy for foraging and general vigor in biocontrol agents (Paul et al., 2022; Rasmussen et al., 2018; Wong et al., 2021) as well as determining their daily activity patterns (Abe et al., 2014; Ogino et al., 2019). Each monitor can hold 32 glass tubes, and automatically and continuously counts how many times the occupant of each tube crosses a set of infrared beams at the middle as long as the monitor is operating. We set the monitors to tabulate and record the number of beam crosses over five-minute time intervals for the duration of both experiments. Age-synchronized parasitoids from rearing (<48 h old) were added to glass tubes (12.5 cm long, 2.5 cm diameter) which had a drop of honey (*ad libitum*) near each end and were capped with firm cotton plugs (narrow Flugs, Diamed Lab Supplies Inc., Mississauga, Canada). The parasitoid loading process was split into blocks of 8 tubes for each of the four light treatments, which were loaded on the same or consecutive days as soon as enough parasitoids emerged to fill a full block in all four treatments. In each of the two temporal replicates, every light treatment received 16 females and 16 males (four of each per loading block), for a total of 128 parasitoids (64 females and 64 males). This amounted to a grand total of 256 parasitoids (128 females and 128 males) in the experiment. The parasitoids were kept in the activity monitors for 12 days, counting from each block’s loading date. This produced 11 full days of data, since the incomplete first day, when the parasitoids were put into the monitors, was omitted. This duration was meant to encompass the parasitoid’s adult lifespan (Shalaby and Rabasse, 1979). Most, though not all, of the parasitoids died—or at least stopped moving entirely—before the end of the experiment, which provided an opportunity to examine whether lighting treatments affected the active lifespan. Upon removal, the tubes were checked to verify whether the parasitoids were alive or dead, to provide additional validation for the activity monitors’ readings. In the photoperiod experiment, 40 of 256 parasitoids were visually confirmed as still alive the day after the experiment ended, and in the spectral quality experiment 75 of 256 were still alive.

### 2.5 Experiment 2: The effect of spectral quality of day-length extensions on aphid reproduction and parasitoid activity

In the spectral quality experiments, we sought to determine the effects on greenhouse insects of the spectral quality of light used to artificially extend the day. We applied four light treatments: three “day extension” treatments of different spectral qualities and one control. The spectral qualities used consisted of the same BSW treatment as described for the photoperiod experiments, as well as two more consisting entirely of narrowband LEDs (Figure 1): a red and blue treatment (RB) and a red, green, and blue treatment (RGB). The RB treatment had peaks at 450 nm (blue), 620 nm (red), and 660 nm (red). The RGB had peaks at 450 nm, 530 nm (green), 620 nm, and 660 nm. The BSW treatment was, as before, chosen to include the wavelengths that are found in sunlight (as much as possible) and to match the photoperiod experiment. The RB treatment was chosen because red and blue LEDs are energetically efficient and photosynthetically efficient, are commercially available for use in greenhouses, and their effects on plants have been widely studied (Cocetta et al., 2017; Massa et al., 2008). The RGB treatment contains red and blue for the same reasons as the RB treatment, and the addition of green reflects more recent research indicating that green light may be valuable for its ability to penetrate more deeply into dense canopies (Smith et al., 2017). Replacing some of the red with green also increased the proportion of shorter wavelengths that are within the insects’ peak spectral sensitivity range (Briscoe and Chittka, 2001).

The mean intensities for the light settings were between 82.2 and 90.5 μmol photons m^-2^ s^-1^ at insect height, including all wavelengths from 250–850 nm. More details are shown in the appendix (Table A.1). All light treatments turned on at 04:00 at the BSW setting as described for the photoperiod experiments, and remained on for 12 hours to produce the “baseline day,” approximating a day where light is predominantly provided by broad-spectrum natural light. At 16:00, the lamps changed to either the RB setting (“RB extension”) or the RGB setting (“RGB extension”), or remained on the BSW setting (“BSW extension”), or turned off (“no extension”, i.e. control). This was designed to simulate a transition to evening where light is predominantly provided by artificial lighting, or where darkness falls. All lights that remained on for the “extension” period turned off at 22:00, producing days of either 18L:6D for the three extension treatments or 12L:12D in the non-extended treatment. A 12-hour baseline day is close to the median day length at the latitude where the research was conducted (National Research Council Canada, 2020), and a 6-hour extension was chosen to be long enough for responses to the altered spectrum to be detectable, while also falling within the range selected for the photoperiod experiment.

Aphid reproduction and parasitoid locomotor activity under the different spectral quality treatments were measured in the same way as described in section 2.4.

### 2.6 Statistical analysis

All statistical analyses were conducted in R version 4.1.2 (2021-11-01) (R Core Team, 2021).

#### Aphid reproduction

Aphid peak fecundity was measured as the total number of nymphs produced by the three aphids on each leaf over the 18-day experimental period. Some mother aphids died before the experiment concluded. For a few leaves, a nymph was missed in counting, reached adulthood, and began to reproduce; these leaves were omitted from the statistical analysis. For the photoperiod experiment, one leaf was omitted from the 14-hour photoperiod treatment, leaving a total of 79 leaves. In the spectral quality experiment, two leaves were omitted from the BSW extension treatment, leaving a total of 78 leaves. For the second replicate of both the photoperiod and spectral quality experiments, where the final aphid count was on the 19^th^ rather than 18^th^ day, an estimate for the number of nymphs produced by day 18 was obtained by assuming an equal number were produced each day since the previous count and interpolating a value. Differences between light treatments were tested with a Kruskal-Wallis test due to the residuals violating the normality assumption for a linear model.

#### Parasitoid locomotor activity and lifespan

Analysis of activity monitor data was performed using a custom R script that built on the *rethomics* suite of R packages (Geissmann et al., 2019). Parasitoid locomotor activity was aggregated by experiment day, with time 0 in each day defined as the time the lights turned on. For each parasitoid, data were only averaged across days for which at least one full day of data was available (see below). Key parameters were (a) total daily activity, defined as the mean number of logged beam crosses per parasitoid per day, averaged across all days; (b) time spent active per day, defined as the mean number of 5-minute intervals over each 24 hours in which the parasitoid moved at least once, averaged across all days; and (c) peak activity time, defined as the hour after the lights turned on in which the parasitoid logged the most beam crosses, averaged across all days.

If, during the experiment, a parasitoid ceased moving and was never detected crossing the beams again, data after that point (which consisted of constant zero values) was omitted, and the time of movement end was logged for survival analysis. The incomplete final day of life was omitted from the activity analysis. Given that this species is diurnal, the earliest time the lights turned off on the final day was considered to be the end of the experiment for the survival analysis to avoid treating resting parasitoids as dead. A total of 46 of 256 parasitoids in the photoperiod experiment and 35 of 256 in the spectral quality experiments were found to have yielded less than one full day of movement data and were omitted entirely from the activity parameter analyses. One female in the BSW extension treatment escaped prior to the experiment start time and was excluded from all analyses. The lifespan survival analysis omitted only parasitoids that died or escaped during the incomplete, omitted first day before the experiment began. A full breakdown of parasitoids included and omitted in each analysis is included in Table A.2.

The activity parameters of daily beam crosses and daily active bins were analyzed using linear mixed models constructed with the *lme4* package (Bates et al., 2015) using the restricted maximum likelihood (REML) method, with log or square root transformations applied when needed to satisfy assumptions of homoscedasticity and normality of residuals, which were checked visually (see *Results*). Factors considered in the models were the fixed effects of light treatment (photoperiod or extension spectral quality), sex, temporal replicate (which had too few levels to be considered as a random effect without producing a singular fit), and loading block within replicate as a random effect. For the peak activity time models in both experiments, including the random effect of blocks produced a singular fit, so the random block effect was removed and linear models were fit including only the fixed effects, which did not change the model estimates (Pasch et al., 2013). Models were assessed using the package *lmerTest* (Kuznetsova et al., 2017) and *car* (Fox and Weisberg, 2019; Kuznetsova et al., 2017) with a type II ANOVA. Models were simplified through backwards stepwise selection, assessing the significance of parameters within the model using an F-test and removing non-significant factors, starting with interactions (Crawley, 2013). The Satterthwaite approximation (Satterthwaite, 1946) was used to estimate the degrees of freedom for ANOVA tests on mixed models. For the effects of extension treatment on the number of beam crosses, the simplified model was used to analyze effects of light treatment. This was done with post-hoc Tukey pairwise tests on marginal means with the *emmeans* package (Lenth, 2022), using the Kenward-Roger approximation for degrees of freedom. For the effects of photoperiod on peak time, the simplified model was used to conduct Tukey pairwise tests on the means averaged over sex. For the effects of extension on peak time, where the interaction between light treatment and sex was significant, separate models were produced for females and males, and Tukey pairwise tests were conducted on each of these.

Parasitoid survival was approximated by the active lifespan as described above, and analyzed post-hoc with a Cox proportional hazards model using the packages *survival* (Therneau and Grambsch, 2000) and *survminer* (Kassambara et al., 2021) to check whether the light treatments influenced parasitoid active lifespans. For the photoperiod experiment, the survival analysis was stratified by sex in order to satisfy the proportionality assumption of the Cox model (Kleinbaum and Klein, 2012). Post-hoc Tukey pairwise tests were performed on the Cox model to determine the differences between photoperiod treatments. For the spectral quality experiment, the extension treatment violated the proportionality assumption of the Cox model, so a Kruskal-Wallis test of the active lifespan was used instead. Parasitoids that survived until the cut-off time were all assigned an identical rank to approximate right-censoring of the data. A post-hoc Dunn test with a Benjamini-Hochberg correction was used to determine the differences between extension treatments, using the package *dunn.test* (Dinno, 2017).

## 3. Results

### 3.1 Experiment 1: The effect of photoperiod on aphid reproduction and parasitoid activity

#### 3.1.1 Aphid reproduction

Aphid reproduction did not vary with photoperiod treatment (Kruskal-Wallis chi-squared value = 0.162, df = 3, p = 0.98) (Figure 2). The number of offspring produced was similar between treatments, with differences of at most nine nymphs between treatment means, amounting to no more than a 3.7% difference.

**Figure 2.**
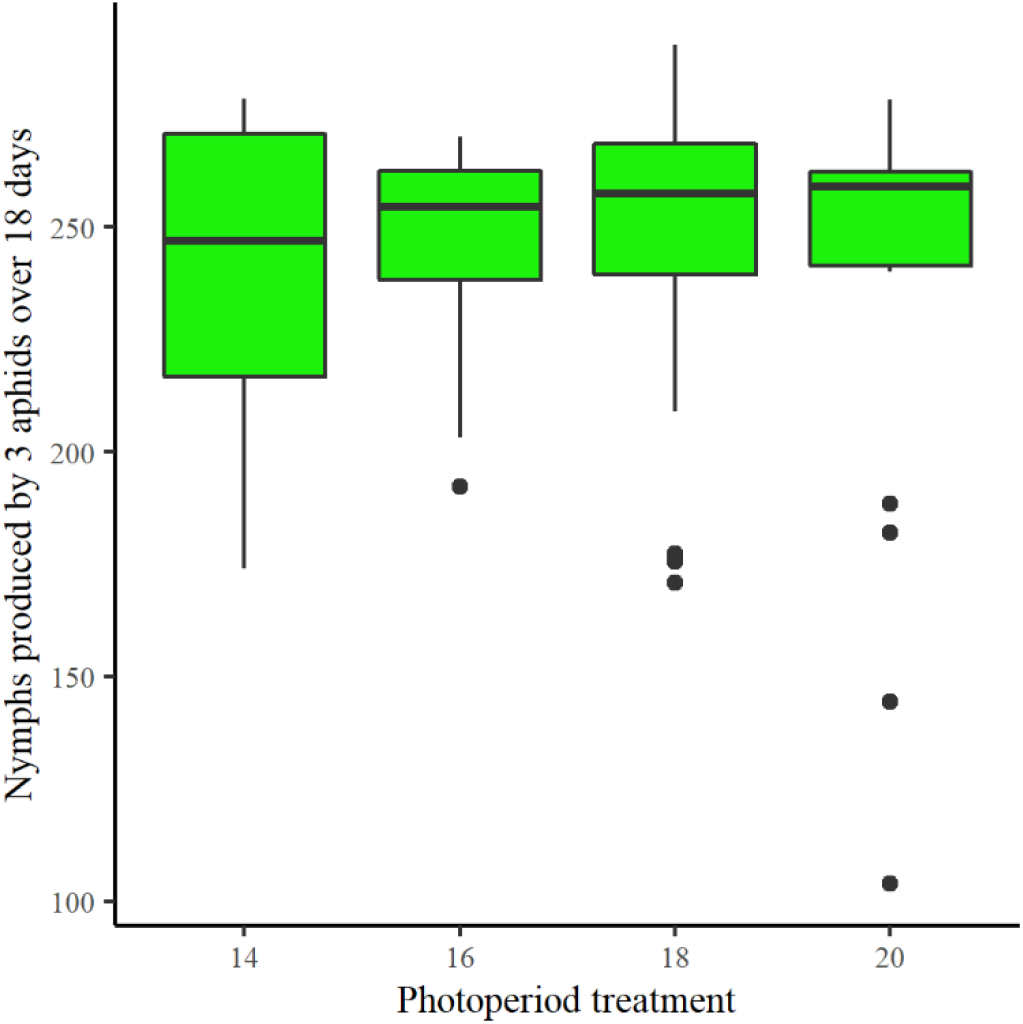
*Myzus persicae* peak fecundity did not vary with photoperiod. Peak reproduction of *M. persicae* was measured under 14L: 10D, 16L:8D, 18L:6D, and 20L:4D photoperiod regimes using a broad-spectrum 5700K white LED plus narrowband LEDs with peaks at 380 nm, 620 nm, and 735 nm (average total photon flux density 87.4 μmol m^-2^ s^-1^). No significant difference was observed between photoperiod treatments (p = 0.98).

#### 3.1.2 Parasitoid locomotor activity and active lifespan

Parasitoids redistributed their activity under photoperiods ranging from 14 to 20 hours without changing how much they moved overall (Figures 3 and 4, Table 1). They clearly displayed diurnal activity patterns, starting to move promptly after the lights turned on, stopping when they turned off, and moving very little in the dark (Figure 3).

**Figure 3.**
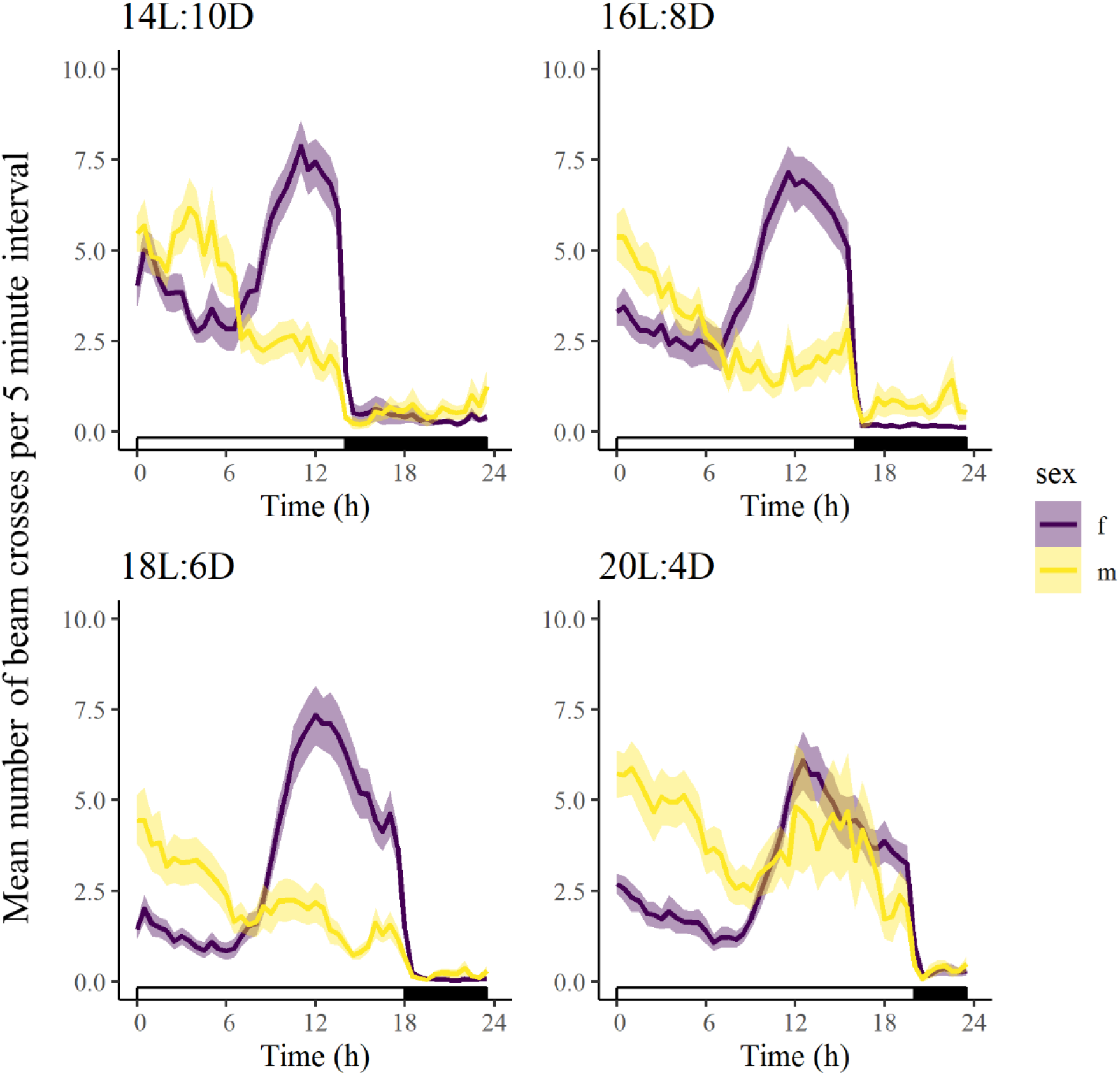
Parasitoids showed clear diurnal activity patterns and adjusted their activity timing to match the photoperiod, and females had later, but more intense, activity peaks than males. Mean (solid lines) and standard error (pale outlines) of locomotor activity (mean number of beam crosses per 5-minute interval) were computed for female and male *Aphidius matricariae* under 14L:10D, 16L:8D, 18L:6D, and 20L:4D light regimes using a broad-spectrum 5700K white LED plus narrowband LEDs with peaks at 380 nm, 620 nm, and 735 nm (average total photon flux density 87.4 μmol m^-2^ s^-1^). Data are averaged across all 11 days in the experiment, though not all individuals survived for the full duration.

**Figure 4.**
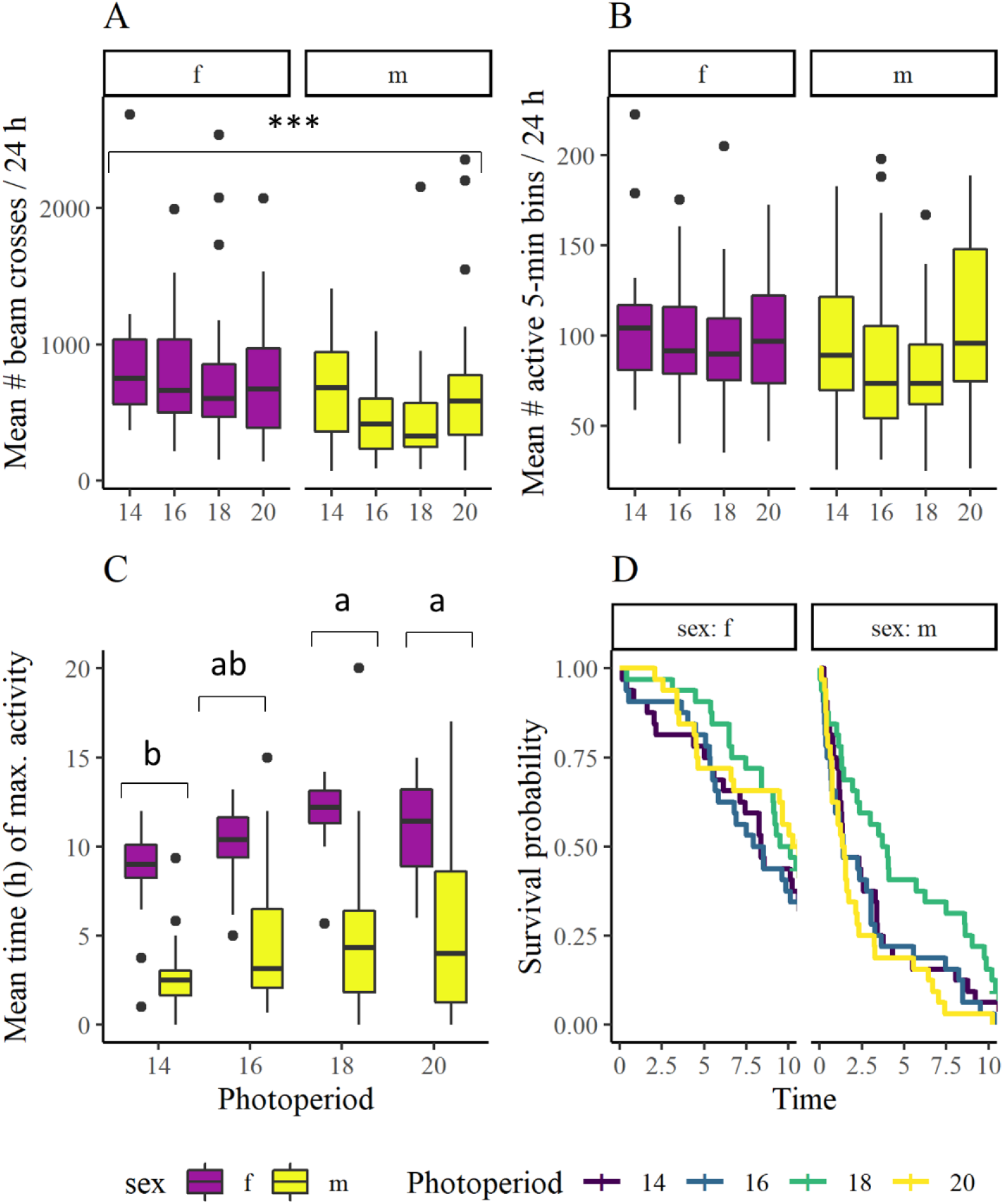
Aphidius matricariae changed the timing, but not the overall amount, of its daily locomotor activity as photoperiod increased from 14 to 20 hours, and had the greatest active lifespan under 18-hour days. Locomotor activity of female and male *A. matricariae,* as a predictor of foraging activity, was examined under 14L: 10D, 16L:8D, 18L:6D, and 20L:4D photoperiod regimes illuminated with a broad-spectrum white LED light over 11 days or until individuals permanently ceased moving. Data displayed include two temporal replicates. The total daily activity (A) was not observed to vary according to the photoperiod (p = 0.12) but was on average 30% higher in females than males (p < 0.0001). The time spent active per day (B) was also not observed to vary significantly either with photoperiod (p = 0.14) or with sex (p = 0.060), though females did spend an average of 40 more minutes active per day than males. Peak activity hour (C) was later in females than males (p < 0.0001) and varied with photoperiod (p < 0.0001); letters represent Tukey pairwise test results averaged across sex and temporal replicate (not shown). Although the overall model was statistically significant (Likelihood ratio test p = 0.035), none of the pairwise comparisons showed a significant difference in active lifespan between the photoperiod treatments (p > 0.05).

**Table 1.** ANOVA results for models fitted on *Aphidius matricariae* activity parameters in photoperiod experiments. Models were specified as response variable ~ photoperiod*sex + replicate + (1|block:replicate) for the mixed model and peak hour ~ photoperiod*sex + replicate for the linear model, and simplified by backwards model selection. Degrees of freedom for the mixed models were obtained using the Satterthwaite approximation. Shaded rows indicate parameters removed during model simplification. Type II ANOVA was performed.

| Response variable (mixed models) | Parameter | Numerator df | Denominator df approximation | F-value | p-value |
| --- | --- | --- | --- | --- | --- |
| Mean daily beam crosses (log transformed) | Sex | 1 | 203.63 | 20.61 | <0.0001 |
|  | Photoperiod | 3 | 199.03 | 1.94 | 0.12 |
|  | Replicate | 1 | 5.71 | 0.15 | 0.71 |
|  | Photoperiod : sex | 3 | 196.48 | 1.32 | 0.27 |
| Mean daily active bins (5-min intervals, square root transformed) | Sex | 1 | 203.08 | 3.59 | 0.060 |
|  | Photoperiod | 3 | 199.00 | 1.85 | 0.14 |
|  | Replicate | 1 | 5.97 | 0.39 | 0.56 |
|  | Photoperiod : sex | 3 | 196.23 | 1.09 | 0.36 |
| Response variable (linear model) | Parameter | df | F-value | p-value |  |
| Mean peak activity hour (hours since lights turned on) | Photoperiod | 3 | 8.91 | <0.0001 |  |
|  | Sex | 1 | 163.42 | <0.0001 |  |
|  | Residuals | 205 |  |  |  |
|  | Replicate | 3 | 3.25 | 0.073 |  |
|  | Photoperiod : sex | 3 | 0.88 | 0.45 |  |

Despite their obvious sensitivity to light, the parasitoids did not change how much they moved overall under different photoperiods (F = 1.96, p = 0.12; Figure 4A) or how much time they spent active each day (F = 1.86, p = 0.14; Figure 4B), suggesting they took more or longer breaks in their locomotor activity during longer days. However, the timing of their period of most intense activity did depend on photoperiod (F = 8.86, p < 0.0001). Parasitoids tended to move their peak activity time later in the day under longer photoperiods, peaking later under 18- and 20-hour days than 14-hour days (p < 0.001 in both cases; Table A3).

The parasitoids’ active lifespan was only marginally affected by the photoperiod and the temporal replicate; the overall model was statistically significant (Likelihood ratio test statistic = 10.57, df = 4, p = 0.032, Table A4) but none of the pairwise models indicated a statistically significant difference between the photoperiod treatments (p > 0.05; Table A-5). Females tended to live longer than males (Figure 2-4D), with females remaining active for, on average, 7.96 (SE 3.18) days and males 3.27 (SE 0.29) days of the possible 11.

There were also several sex differences in activity patterns, though they did not influence the parasitoids’ responses to changing photoperiods. Males reached their activity peak earlier in the day than females (p < 0.0001) while females had later, more intense activity peaks (Figure 3), resulting in higher total daily activity in females (F = 20.50, p < 0.0001) by an average of 30% (Figure 4A). The time spent active per day (Figure 4B), however, did not differ significantly between females and males (F = 3.42, p = 0.06).

A more complete description of the activity models is presented in Table A.3, results of pairwise comparisons in Table A.4, and results of the survival model in Table A.5.

### 3.2 Experiment 2: The effect of spectral quality of day-length extensions on aphid reproduction and parasitoid activity

#### 3.2.1 Aphid reproduction

The number of nymphs produced by aphids did not vary depending on the duration of LED daylength extensions (Kruskal-Wallis chi-squared value = 7.57, df=3, p= 0.056), though the aphids tended to reproduce slightly more under the extended 18-hour days compared to the non-extended 12-hour days (Figure 5).

**Figure 5.**
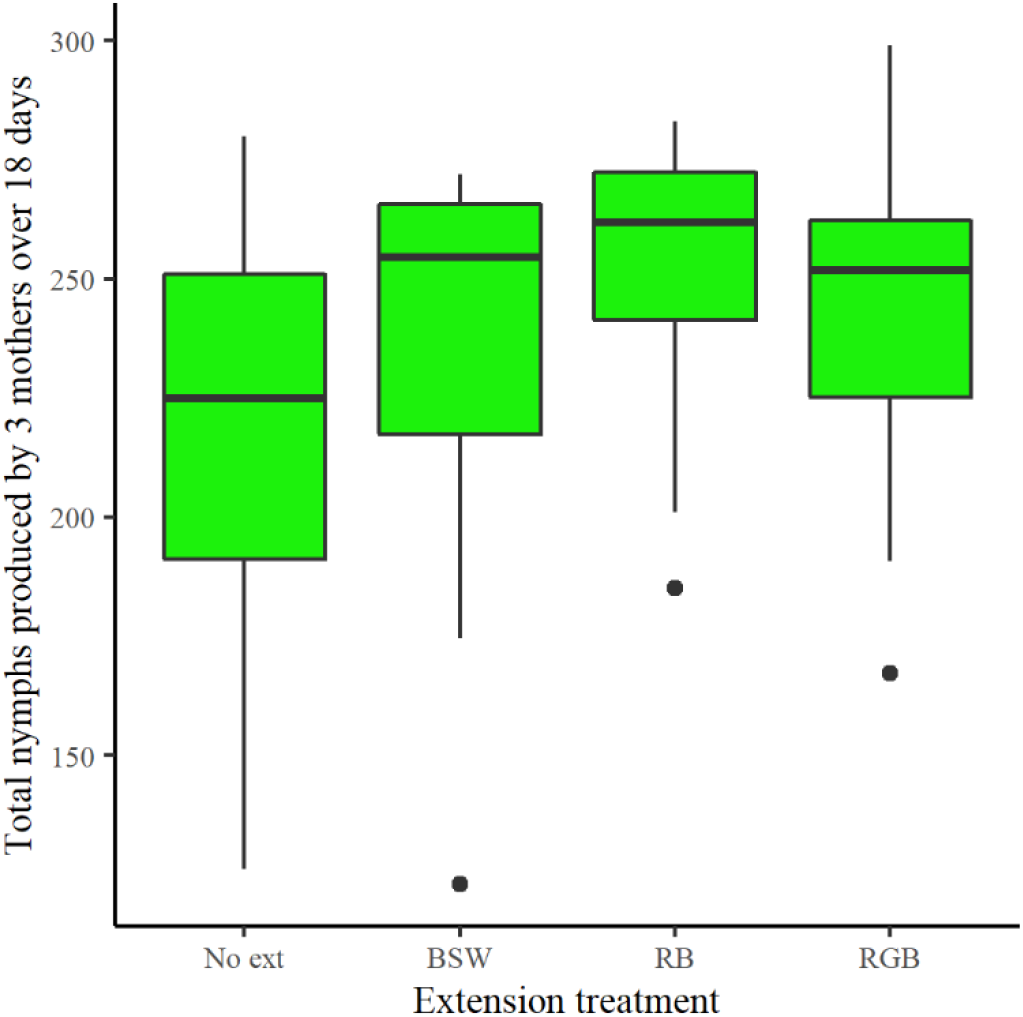
*Myzus persicae* peak fecundity did not vary significantly with LED day extension treatment. Peak reproduction of aphids was studied under 12-hour days (“no extension”) or 12-hour days supplemented with an additional 8 hours of broad-spectrum white (BSW), red and blue light (RB), or red, green, and blue light (RGB). No significant difference was observed between treatments (p = 0.056).

#### 3.2.2 Parasitoid locomotor activity under day extensions of different spectral quality

Parasitoids adjusted their activity timing and duration in response to LED day extensions, some of which was clearly in response to the spectral quality used—and not solely the lengthening of the photoperiod— but they did not change their total amount of activity under any treatment (Figures 6 and 7, Table 2). They displayed the same diurnal patterns as in the photoperiod experiment, remaining active under all spectral qualities used (Figure 6). Daily activity peaks again tended to decrease in intensity as the experiment progressed (Figure A.2).

**Figure 6.**
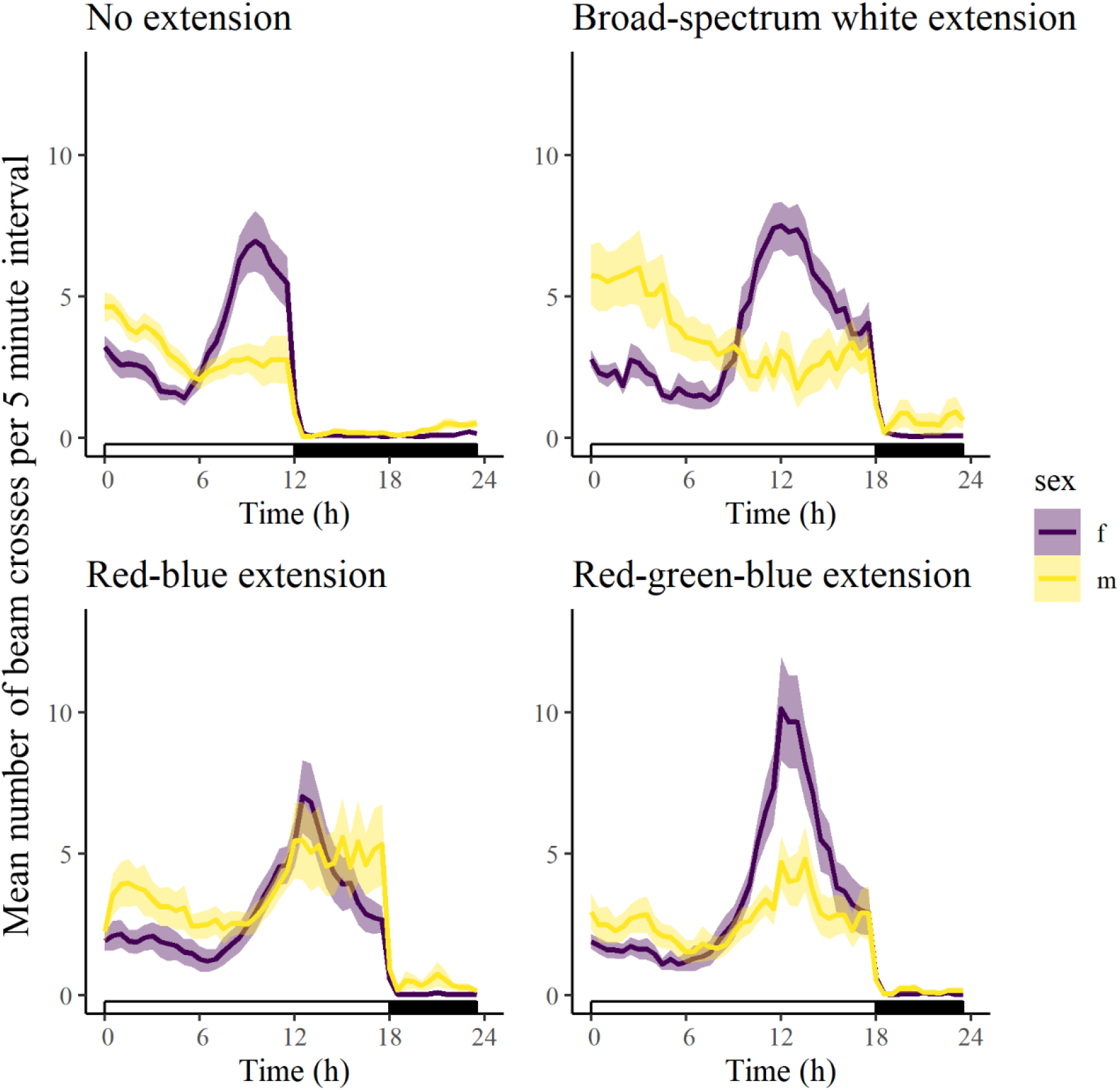
Parasitoids adjusted their daily activity timing under LED supplementation of all spectral qualities tested. Mean (solid lines) and standard error (pale outlines) of activity (mean number of beam crosses per 5-minute interval) over time were computed for female and male *Aphidius matricariae* under 12-hour days (“no extension”) or 12-hour days supplemented with an additional 8 hours of broad-spectrum white (BSW), red and blue light (RB), or red, green, and blue light (RGB). Data are averaged across all 11 days of the experiment, though some parasitoids did not survive that long. Males in both the RB and RGB extensions, and females in the RGB extension, showed especially high activity intensity shortly after the lights switched from the BSW spectrum to the treatment spectrum at 12 hours after the lights turned on.

**Figure 7.**
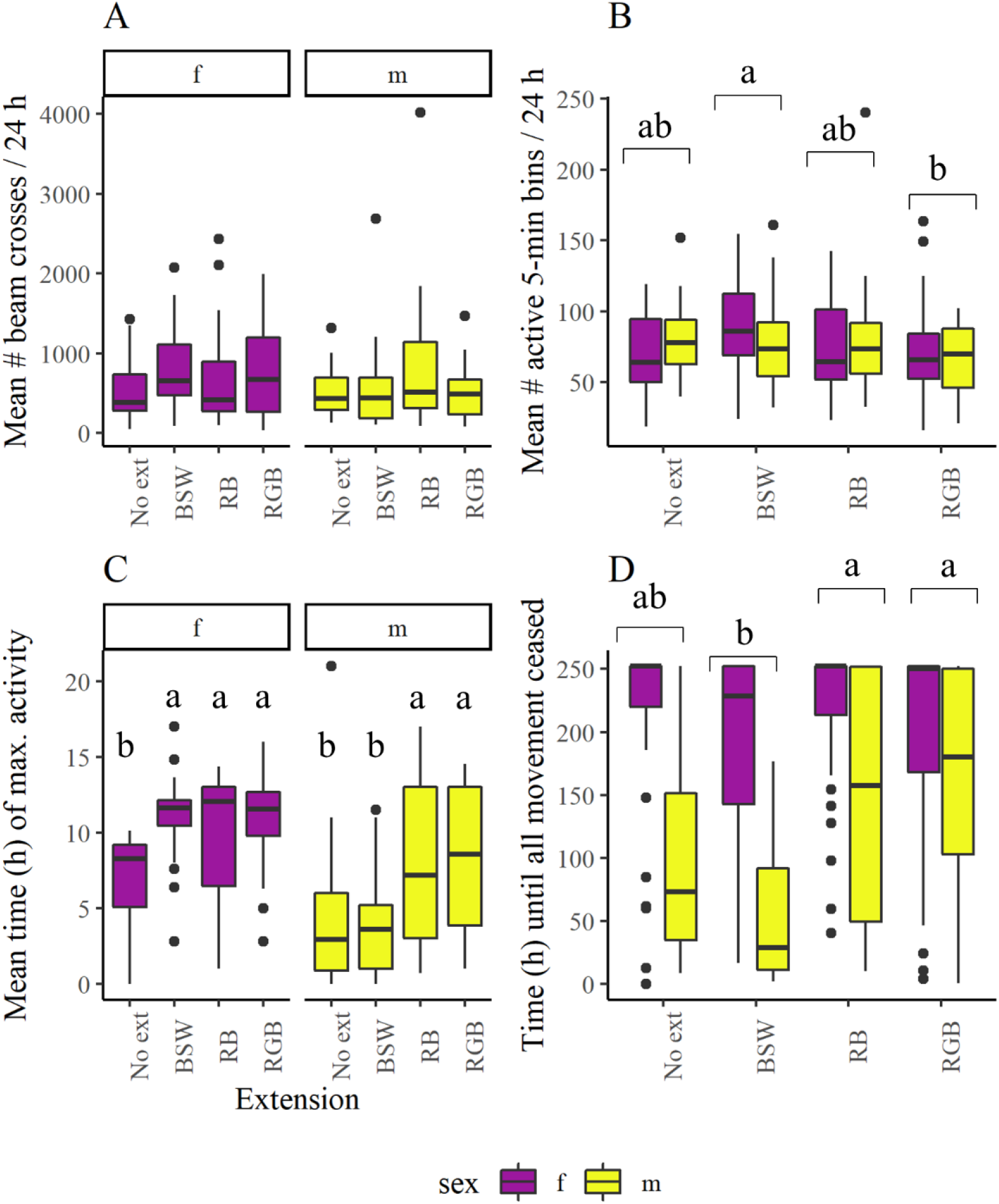
*Aphidius matricariae* changed the timing and duration of its daily locomotor activity under different LED day extension regimes, but not the overall amount. Locomotor activity of female and male *A. matricariae,* as a predictor of foraging activity, was examined under 12-hour days of broad-spectrum white LED light (No ext) or identical 12-hour days supplemented with an additional 8 hours of broad-spectrum white (BSW), red and blue light (RB), or red, green, and blue light (RGB). Data displayed include two temporal replicates. The total daily activity (A) did not vary according to light extension treatment (p = 0.41), but the time spent active (B) did (p = 0.033), with parasitoids in the BSW treatment spending an average of 90 more minutes active than those in the RGB treatment (p = 0.019). Peak activity hour (C) varied with sex (p < 0.001), extension treatment (p < 0.001), and the interaction between them (p = 0.004). Letters in (C) represent results of Tukey pairwise comparisons and apply only within the same sex. Parasitoid active lifespan (D) varied with extension treatment (p = 0.0010), with parasitoids under the RB treatment having the longest active lifespan and those under the BSW treatment having the shortest.

**Table 2.**
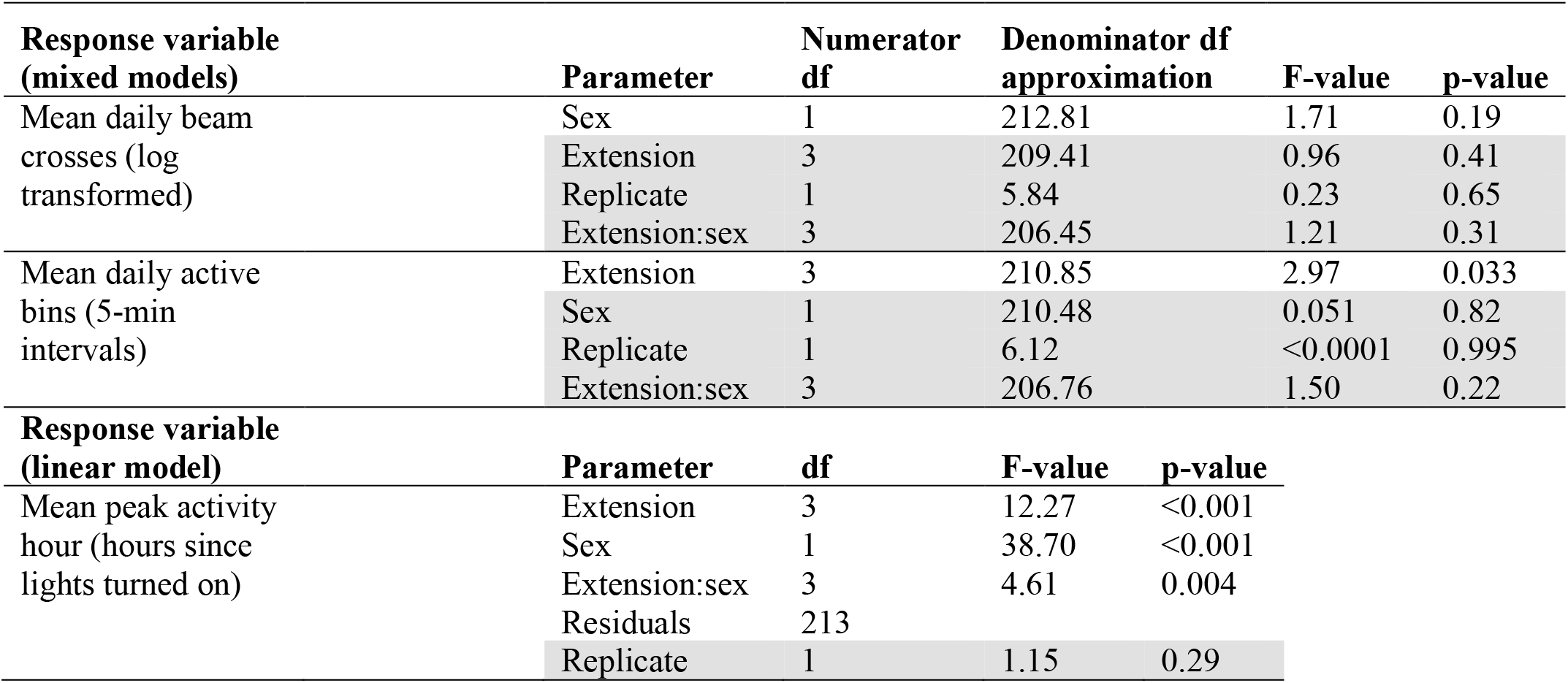
ANOVA results for models fitted on *Aphidius matricariae* activity parameters in spectral quality experiments. Models were specified as response variable ~ extension*sex + replicate + (1|block:replicate) for mixed models and response variable ~ photoperiod*sex + replicate for the linear model, and simplified by backwards model selection. For the mixed models (beam crosses and active bins), degrees of freedom were obtained using the Satterthwaite approximation. Shaded rows indicate parameters removed during model simplification. Type II ANOVA was performed.

While the parasitoids’ total daily activity did not vary with the day extension treatment (F = 0.94, p = 0.41), the time they spent active did (F = 3.03, p = 0.033). Parasitoids spent more time active under the BSW treatment than under the RGB treatment (t = 2.95, p = 0.019) by an average of approximately 90 minutes per day (Table A.6). The timing of their peak activity period during the day also depended on the extension treatments in a sex-dependent manner (F = 4.51, p = 0.004). Females had later activity peaks under days extended with any spectral quality relative to the non-extended treatment by 2.7–4.1 hours on average (Table A.6). Males, on the other hand, peaked later, by 4.0 to 4.6 hours on average, under the RB and RGB treatments compared to the 12-hour non-extended day treatment and the BSW-extension treatment (Table A.6). The males’ peak times under the RB and RGB treatments were closer to those shown in females than in males under either of the BSW-only treatments (Figures 6 and 7C). Thus, the males’ observed peak time response was sensitive to the spectral quality used, while the females’ response depended only on the total photoperiod, regardless of the spectral quality of the light used to produce it. These changes appear to have taken place primarily during the extended period of the day after the lights switched from BSW to the altered spectrum at 12 hours after the lights came on (Figure 6).

The parasitoids’ active lifespan was also affected by the spectral quality (Kruskal-Wallis chi squared = 16.24, df = 3, p = 0.0010). Parasitoids under the RGB and RB treatments survived significantly longer than those under the BSW treatment (p = 0.0019 and p = 0.0027, respectively; Table A-7). None of the extended (18h photoperiod) treatments had a significantly different active lifespan from the shorter-day control (p > 0.05). Females tended to have longer active lifespans than males (Figure 7D), with females remaining detectably active for an average of 8.45 (SE 0.28) days and males 4.85 (SE 0.24) days out of a possible 11, though we were unable to test for a difference. Sex differences were not observed for the other activity parameters, with females and males showing no clear differences in total activity (F = 1.32, p = 0.25) or time spent active (F = 0.049, p = 0.83).

A more complete description of the models is presented in Table A.6, and results of pairwise comparisons in Table A.7.

## 4. Discussion

Our goal in this set of experiments was to determine how extending the day affects aphid reproduction and parasitoid locomotor activity, both through the duration of the extension and the spectral quality used. We found that aphid fecundity was mostly unresponsive to photoperiod and not affected by the spectral quality of daylength extensions. Parasitoids shifted the timing of their locomotor activity to remain active later during days with longer photophases, and showed the most intense activity under RGB-extended days, but were never observed to change their total amount of daily activity. This is, to our knowledge, the first experiment to show how parasitoids modify their locomotor activity over multiple days in response to greenhouse lighting incorporating novel spectral qualities.

### 4.1 Effects of photoperiod and spectral quality on aphid reproduction

We observed no significant effects of photoperiod on aphid fecundity. This is similar to the findings of Joschinski et al. (2015) on *Acyrthosiphon pisum* (Hemiptera: Aphididae), which showed that the direct effect of lengthening the photoperiod was merely a minor increase in fecundity and reproductive lifespan that proved to be insignificant at the population level. While some mechanisms have been proposed for photoperiod to affect aphid fecundity or feeding, such as disruption of circadian rhythms (Joschinski et al., 2015) or changes in phloem osmotic pressure (Cull and van Emden, 1977; Nalam et al., 2021), any effects of photoperiod that may have occurred in our experiments were too subtle to detect. Our results support the idea that the most useful effects of manipulating spectral quality on aphids will be in keeping aphids away from host plants, by disrupting their navigation and host-finding or attracting them for trapping (Ben-Yakir et al., 2012; Johansen et al., 2011; Martini et al., 2020; Shimoda and Honda, 2013). However, we might expect to observe stronger impacts of both photoperiod and spectral quality on aphids on whole plants, where they will be susceptible to the bottom-up effects of light on the system (Vänninen et al., 2010); we examined this in later research (Fraser, 2022).

### 4.2 Effects of photoperiod and spectral quality on parasitoid locomotor activity

#### 4.2.1 Total parasitoid locomotor activity

Contrary to our predictions, parasitoids did not increase their total activity as day lengths increased, or change it under day extensions of different spectra, though they did redistribute it to different times of the lit period of the day. The total activity level was probably determined by a combination of other elements including nutrition, age, sex, genetics, and other physiological factors (Pompanon et al., 1999; Varennes et al., 2016). Since we observed the parasitoids distributing the same active time and same amount of movement over increasingly long photoperiods, they must have spent more or longer periods stationary during the photophase under longer days. A similar tendency to spread the same amount of locomotor activity over a longer day was observed in *Aphidius gifuensis* (Hymenoptera: Braconidae) (Abe et al., 2014). A different response manifested in our spectral quality experiment, where the spectral quality of the light used did influence the total duration that *A. matricariae* spent active—though again without changing the total amount of locomotor activity per day.

The fact that the parasitoids did not change their total daily activity in response to the light conditions might be because they were already at their maximum daily spontaneous activity level under all the light treatments we tested. Animals will not necessarily increase their daily locomotor activity beyond the minimum amount necessary for survival (Herbers, 1981), and when they increase how fast they move or how long they are active, they must account for trade-offs such as an increased risk of being preyed upon (Werner and Anholt, 1993). Given that the parasitoids were fed *ad libitum,* they would not have been limited in their locomotor activity by lack of a sugar source the way unfed parasitoids are (Siekmann et al., 2004; Varennes et al., 2016). It is possible that more extreme (e.g. shorter-day) light treatments than we tested could have curtailed their activity. Alternatively, *A. matricariae* could be comparatively insensitive to light conditions that other parasitoids might have responded to, and in that case, a species-specific approach would be needed to understand the effects of LED illumination on biocontrol agents.

We found that female *A. matricariae* were more active than males under the photoperiod experiment, which is similar to previous findings in *A. gifuensis* (Abe et al., 2014). We also found male activity consistently peaked earlier in the day than female activity, similarly to some other parasitoid species (Bertossa et al., 2013; Ndoutoume-Ndong et al., 2006; Pompanon et al., 1999). More notably, we found that males, but not females, were sensitive to the spectral quality when determining their peak locomotor activity time under extended days. This asymmetric effect on the two sexes might impact mating behaviors, as males typically emerge early in the day to mate and disperse (Bourdais and Hance, 2019); this could be a topic for future research.

#### 4.2.2 Parasitoid activity timing

We found that males adjusted their peak activity time in response to spectral quality, but females did not, and that both females and males spent more time active per day under the BSW treatment compared to the RGB treatment. These changes appear to have occurred specifically during the extension period. The females’ peak activity times in all three extended treatments were also close to the 12-hour mark, which is when the light regime switched over from the “baseline day” to the extension treatment. The RGB-treatment females, in particular, showed a sharp activity spike during the RGB-extended portion of the day, implying a bout of intense activity following the change in spectral quality, followed by “compensation” for the increased energy expenditure. We hypothesize that this result could have been due to parasitoids having more intense activity when lights changed to a treatment with shorter wavelengths (e.g., substituting red or far-red photons with green ones) that they perceived as “brighter” (Barbosa and Frongillo, 1977; Cochard et al., 2017; Peitsch et al., 1992), although this would need to be validated by further research.

#### 4.2.3 Parasitoid longevity

Parasitoid active lifespan was affected by the light treatments. In particular, parasitoids under the RB- and RGB-extended photoperiods showed longer adult lifespans than those under the 18-hour BSW treatment, though from this experiment we cannot determine why. Reported longevity for *A. matricariae* at 20–21°*C* ranges from roughly 6–7 days (Giri et al., 1982; Pourtaghi et al., 2016; Rashki et al., 2020) to 14 days (Shalaby and Rabasse, 1979), with no significant differences between females and males, depending on the food they are provided. Under our colony rearing conditions, adults frequently lived for two weeks or more (J. Fraser, personal observation). In the experiments, the mean active lifespan was about 8 days for females and about 3-5 for males of the 11 day duration, so the experimental conditions may have shortened the adult lifespan for males in particular.

### 4.3 Conclusions

Our most notable finding was that parasitoid behaviour was highly flexible in response to drastic changes in lighting: *A. matricariae* was able to quickly adjust its total daily activity levels to photoperiods and light spectra very different from those under which it was reared. If this finding extends to other parasitoid species, it is promising for the prospect of releasing parasitoids in greenhouses with lighting conditions very different from parasitoid rearing facilities and from the insects’ native environments. The fact that the duration and peak timing of activity of parasitoids did nonetheless shift under different day lengths and spectra is also notable, and may have implications for their efficacy if the timing of parasitoid foraging activity interacts with changes in host behavior or physiology that influence their susceptibility to parasitoids (Johansen et al., 2011). The next step in determining the effects of lighting conditions on insect biocontrol in greenhouses will be to test how these changes in natural enemy behaviour translate to differences (or lack thereof) in their efficacy in suppressing host populations and protecting plants.

## Acknowledgements

We would like to thank Clarissa Capko and Jason Thiessen for support in running experiments; Yonathan Uriel for supplying data and insect rearing methodologies; Ricardo Sarte, Paige Boegarts, Denika Joiner, Autumn White, and Angela Oscienny for greenhouse support.

