## Supplementary Information: Tables A.1-A.7, Figures A.1-A.2 for "The effects of LED daylength extensions on the fecundity of the pest aphid *Myzus persicae* and the daily activity patterns of its parasitoid, *Aphidius matricariae*"

**Table A.1 Spectral composition of light treatments used in growth chamber experiments.** The growth chambers ("GC") were numbered 1–4, and treatments were randomly assigned among them for two temporal replicates. The BSW treatment was used in all growth chambers in both sets of experiments. Readings are averaged across 8 positions within each growth chamber, at 64.5 cm below the lamp surface (13.6 cm above the rack), which was the insects' approximate level during the experiments. Measurements were taken using an Optimum SRI-2000 handheld spectrophotometer (Optimum OptoElectronics Corp, Hsinchu, Taiwan).

| | **Mean total photon flux density (μmol $m^{-2}$ $s^{-1}$) by wavelength range** | | | | | |
|---|---|---|---|---|---|---|
| **Total photon flux (μmol $m^{-2}$ $s^{-1}$)** | **UV (<400nm)** | **Blue (400-499.5nm)** | **Green (500-599.5nm)** | **Red (600-699.5nm)** | **Far-red (>=700nm)** | **All wave-lengths** |
| **RB** | **0.0** | **18.3** | **0.5** | **69.9** | **0.6** | **89.2** |
| GC1 | 0.0 | 18.1 | 0.5 | 68.8 | 0.3 | 87.8 |
| GC2 | 0.0 | 18.4 | 0.5 | 70.9 | 0.7 | 90.5 |
| **RGB** | **0.0** | **19.2** | **19.8** | **44.1** | **0.2** | **83.3** |
| GC1 | 0.0 | 18.7 | 18.3 | 45.1 | 0.1 | 82.2 |
| GC3 | 0.0 | 19.7 | 21.4 | 43.1 | 0.3 | 84.5 |
| **BSW** | **1.5** | **11.9** | **25.9** | **22.5** | **25.6** | **87.4** |
| GC1 | 1.4 | 11.7 | 26.0 | 22.3 | 25.0 | 86.5 |
| GC2 | 1.6 | 12.3 | 26.3 | 23.4 | 26.2 | 89.8 |
| GC3 | 1.5 | 12.0 | 25.6 | 22.7 | 25.3 | 87.1 |
| GC4 | 1.6 | 11.6 | 25.5 | 21.9 | 26.1 | 86.7 |

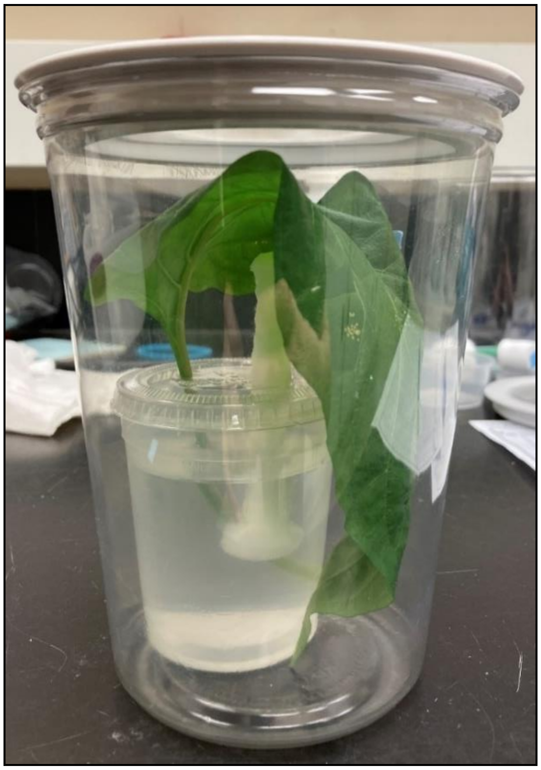

**Figure A.1. Example of containers used in aphid growth chamber experiments.** Each container contained a mature leaf from a 58- or 59-day-old pepper plant and three 6-day-old *Myzus persicae* adults. Leaves were placed in a sealed water container and the container had a mesh lid for ventilation and transparent sides to allow light transmission.

**Table A.2 Summary of treatment numbers in parasitoid experiments.** Parasitoids (*Aphidius matricariae*) with at least one complete day of data in the growth chamber experiments were included in the activity analyses. For each experiment, a total of 64 individuals, comprising 32 females and 32 males, were placed into activity monitors per light treatment for a total of 256 wasps split evenly into two temporal blocks of 128. The wasps omitted from the activity and survival analyses died on or before the first day of the experiment, or escaped (n = 1 female in the BSW treatment).

| | | **Daily activity analysis** | | | **Active lifespan analysis** | | |
|---|---|---|---|---|---|---|---|
| **Experiment** | **Treatment** | **Females** | **Males** | **Total** | **Females** | **Males** | **Total** |
| Growth chamber photoperiod experiment | 14L:10D | 29 | 24 | 53 | 32 | 32 | 64 |
| | 16L:8D | 29 | 19 | 48 | 32 | 32 | 64 |
| | 18L:6D | 31 | 26 | 57 | 32 | 32 | 64 |
| | 20L:4D | 32 | 20 | 52 | 32 | 32 | 64 |
| | Total | 121 | 89 | 210 | 128 | 128 | 256 |
| Growth chamber spectral quality experiment | No extension | 29 | 30 | 59 | 31 | 32 | 63 |
| | BSW | 30 | 16 | 46 | 31 | 31 | 62 |
| | RB | 25 | 32 | 57 | 32 | 32 | 64 |
| | RGB | 29 | 30 | 59 | 32 | 31 | 63 |
| | Total | 113 | 108 | 221 | 126 | 126 | 252 |

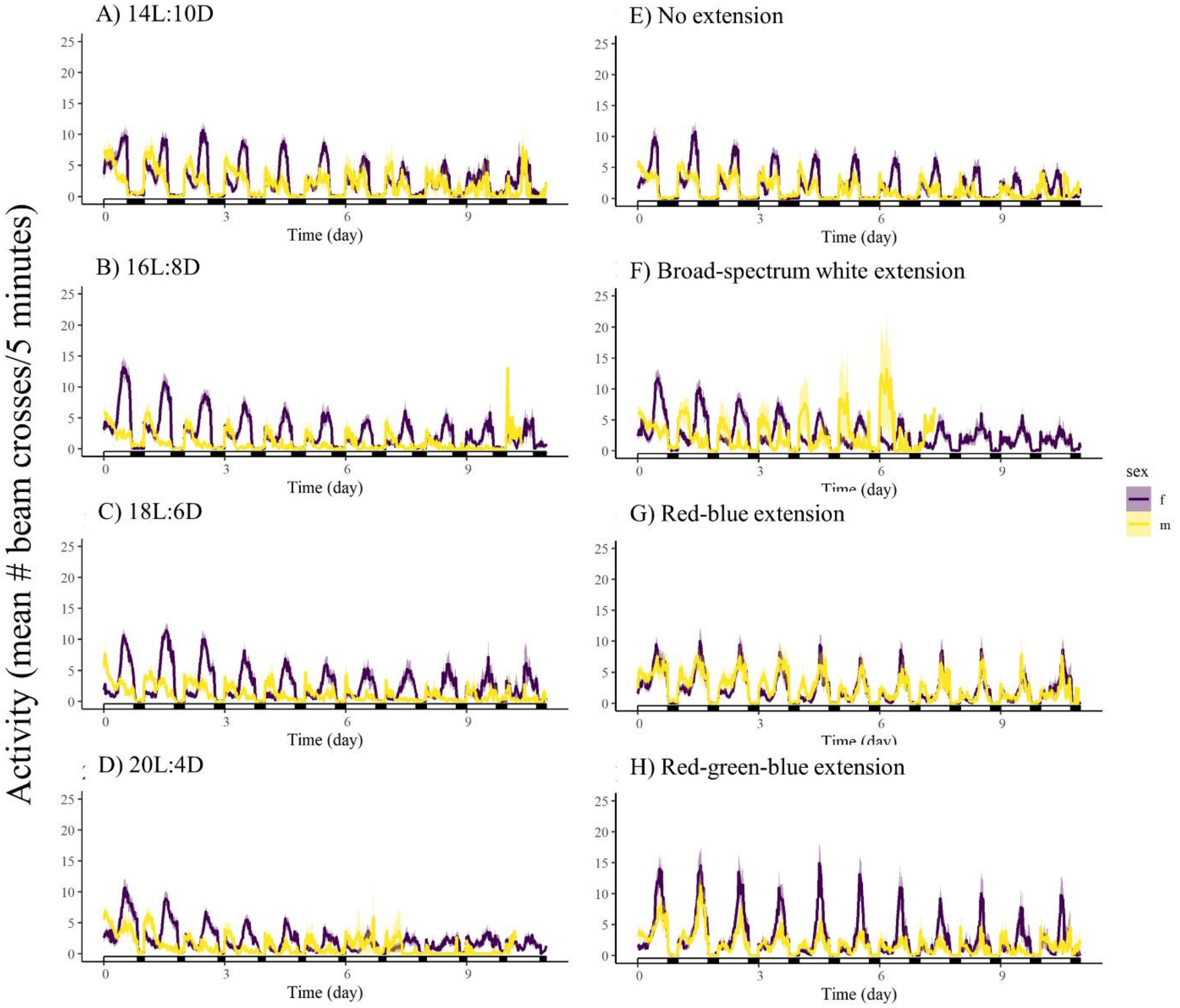


**Figure A.2. Parasitoid activity patterns remained similar throughout the experiments, though activity intensity decreased slightly over time.** Mean (solid lines) and standard error (pale outlines) of locomotor activity over time were computed for female and male *Aphidius matricariae* under (A) 14L:10D, (B) 16L:8D, (C) 18L:6D, and (D) 20L:4D light regimes or a separate experiment with (E) 12-hour days ("no extension") or 12-hour days supplemented with an additional 8 hours of (F) broad-spectrum white, (G) red and blue light, or (H) red, green, and blue light (right). Not all individuals survived the full 11 days, so the means represent the individuals alive at a given time point. In cases where one group is not shown for the full duration of the experiment, this is because all individuals ceased moving before the experiment concluded.

**Table A.3. Summaries of parasitoid activity models in the photoperiod experiment.** Estimates and standard deviations obtained by fitting linear mixed models of wasp (*Aphidius matricariae*) activity parameters in photoperiod experiment are displayed. Mixed models (beam crosses and active bins) were specified as response variable ~ photoperiod*sex + replicate + (1|block:replicate) and the linear model was fitted as mean peak activity hour ~ photoperiod*sex + replicate. Grey-shaded rows indicate factors that were removed in backwards model selection.

| Response variable | Parameter | Estimate of fixed effect | Standard error |
|---|---|---|---|
| Mean daily beam crosses (log transformed) | Intercept | 6.50 | 0.06 |
| | Sex = male | -0.43 | 0.094 |
| | Photoperiod = 16 | -0.21 | 0.13 |
| | Photoperiod = 18 | -0.30 | 0.13 |
| | Photoperiod = 20 | -0.14 | 0.13 |
| | Replicate = second | 0.042 | 0.11 |
| | Photoperiod=16 : sex = male | -0.23 | 0.27 |
| | Photoperiod=18 : sex = male | -0.31 | 0.26 |
| | Photoperiod=20 : sex = male | 0.16 | 0.27 |
| | **Parameter** | **Variance** | **Standard deviation** |
| | Random effect of block within replicate | 0.0017 | 0.042 |
| | Residual | 0.46 | 0.68 |
| **Response variable** | **Parameter** | **Estimate of fixed effect** | **Standard error** |
| Mean daily active bins (5-minute intervals, square root transformed) | Intercept | 9.83 | 0.20 |
| | Sex = male | -0.50 | 0.26 |
| | Photoperiod =16 | -0.41 | 0.37 |
| | Photoperiod =18 | -0.74 | 0.36 |
| | Photoperiod =20 | -0.048 | 0.37 |
| | Replicate = second | -0.22 | 0.35 |
| | Photoperiod = 16 : sex = male | -0.15 | 0.76 |
| | Photoperiod = 18 : sex = male | -0.49 | 0.72 |
| | Photoperiod = 20 : sex = male | 0.80 | 0.74 |
| | **Parameter** | **Variance** | **Standard deviation** |
| | Random effect of block within replicate (intercept) | 0.084 | 0.29 |
| | Residual | 3.56 | 1.89 |
| **Response variable** | **Parameter** | **Estimate of fixed effect** | **Standard error** |
| Mean peak activity hour (hours since lights turned on) | Intercept | 8.67 | 0.49 |
| | Photoperiod = 16 | 1.62 | 0.65 |

| | | | |
|---|---|---|---|
| | Photoperiod = 18 | 2.92 | 0.62 |
| | Photoperiod = 20 | 2.68 | 0.64 |
| | Sex = male | -5.83 | 0.46 |
| | Replicate = second | 0.81 | 0.45 |
| | Photoperiod = 16 : sex = male | 0.66 | 1.31 |
| | Photoperiod = 18 : sex = male | -0.78 | 1.24 |
| | Photoperiod = 20 : sex = male | 1.15 | 1.29 |
| | Multiple $R^2$=0.49 | | Adjusted $R^2$=0.48 |

**Table A.4. Results of pairwise comparisons on parasitoid (*Aphidius matricariae*) activity parameters in the photoperiod experiment.** Tukey pairwise test results on the no-interaction model were specified as mean peak activity hour ~ photoperiod + sex, with means averaged over levels of sex.

| **Response variable** | **Treatment (photoperiod) comparison** | **Estimate (Tukey pairwise tests)** | **SE** | **t-value** | **p- value** |
|---|---|---|---|---|---|
| Mean peak activity hour | 16 – 14 | 1.62 | 0.65 | 2.49 | 0.065 |
| | 18 – 14 | 2.92 | 0.63 | 4.69 | <0.001 |
| | 20 – 14 | 2.68 | 0.64 | 4.20 | <0.001 |
| | 18 – 16 | 1.30 | 0.64 | 2.03 | 0.18 |
| | 20 – 16 | 1.06 | 0.65 | 1.63 | 0.37 |
| | 20 – 18 | -0.24 | 0.63 | -0.379 | 0.98 |

**Table A.5. Summary of survival models in the photoperiod experiment.** Cox proportional hazards model and post-hoc Tukey pairwise comparisons were performed for survival ~ photoperiod + replicate, stratified on sex, for parasitoids (*Aphidius matricariae*) in the photoperiod experiment. The overall model was significant (Likelihood ratio test statistic 10.38, df = 4, p = 0.035).

| **Parameter** | **Regression coefficient** | **Hazard ratio** | **Standard error of regression coefficient** | **Wald statistic Z-value** | **p-value** |
|---|---|---|---|---|---|
| Photoperiod = 16 | 0.12 | 1.13 | 0.20 | 0.63 | 0.53 |
| Photoperiod = 18 | -0.40 | 0.67 | 0.20 | -1.94 | 0.052 |
| Photoperiod = 20 | 0.049 | 1.05 | 0.20 | 0.24 | 0.81 |
| Replicate = second | 0.19 | 1.20 | 0.14 | 1.30 | 0.19 |
| **Treatment (photoperiod) Comparison** | | **Estimate (Tukey pairwise tests)** | **SE** | **t-value** | **p- value** |
| 16 h –14 h | | 0.12 | 0.20 | 0.63 | 0.92 |
| 18 h –14 h | | -0.40 | 0.20 | -1.94 | 0.21 |
| 20 h –14 h | | -0.049 | 0.20 | 0.24 | 0.995 |
| 18 h –16 h | | -0.52 | 0.20 | -2.56 | 0.052 |
| 20 h –16 h | | -0.074 | 0.20 | -0.37 | 0.98 |
| 20 h –18 h | | 0.45 | 0.21 | 2.11 | 0.15 |

**Table A.6. Summaries of parasitoid activity models in the spectral quality experiment.** Estimates and standard deviations obtained by fitting linear or linear mixed models of wasp (*Aphidius matricariae*) activity parameters in spectral quality experiments are displayed. Models were specified as response variable ~ extension*sex + replicate + (1|block:replicate). Mean peak activity hour was fitted as a linear model specified as mean peak hour ~ extension*sex + replicate. Grey-shaded rows indicate factors that were removed in backwards model selection.

| **Response variable** | **Parameter** | **Estimate of fixed effect** | **Standard error** |
|---|---|---|---|
| Mean daily beam crosses (log transformed) | Intercept | 6.24 | 0.093 |
| | Sex = male | -0.14 | 0.11 |
| | Extension = none | -0.23 | 0.16 |
| | Extension = RB | -0.016 | 0.16 |
| | Extension = RGB | -0.12 | 0.16 |
| | Replicate = second | -0.081 | 0.17 |
| | Extension = none : sex = male | 0.46 | 0.33 |
| | Extension = RB : sex = m | 0.52 | 0.33 |
| | Extension = RGB : sex = male | 0.15 | 0.33 |
| | **Parameter** | **Variance** | **Standard deviation** |
| | Random effect of block within replicate | 0.026 | 0.16 |
| | Residual | 0.65 | 0.81 |
| **Response variable** | **Parameter** | **Estimate of fixed effect** | **Standard error** |
| Mean daily active bins (5-minute intervals) | Intercept | 87.37 | 5.04 |
| | Extension = none | -12.65 | 6.23 |
| | Extension = RB | -10.16 | 6.28 |
| | Extension = RGB | -18.36 | 6.23 |
| | Sex = male | 0.97 | 4.33 |
| | Replicate = second | -0.037 | 6.16 |
| | Extension = none : sex = male | 21.3 | 12.84 |
| | Extension = RB : sex = m | 16.5 | 12.95 |
| | Extension = RGB : sex = male | 1.78 | 12.84 |
| | **Parameter** | **Variance** | **Standard deviation** |
| | Random effect of block within replicate (intercept) | 28.48 | 5.34 |
| | Residual | 1002.15 | 31.66 |
| **Response variable** | **Parameter** | **Estimate of fixed effect** | **Standard error** |
| Mean peak activity hour (hours since lights turned on) | Intercept | 11.16 | 0.71 |
| | Extension = none | -4.11 | 1.02 |
| | Extension = RB | -1.37 | 0.99 |
| | Extension = RGB | -0.34 | 1.01 |
| | Sex = male | -7.24 | 1.21 |

| | | | |
|---|---|---|---|
| | Extension = none : sex = male | 4.19 | 1.58 |
| | Extension = RB : sex = m | 5.49 | 1.60 |
| | Extension = RGB : sex = male | 4.95 | 1.58 |
| | Replicate = second | -0.57 | 0.53 |
| | Multiple $R^2$ = 0.30 | Adjusted $R^2$ = 0.28 | |

**Table A.7. Results of pairwise comparisons on parasitoid activity parameters in the photoperiod experiment.** Tukey pairwise tests were performed on the simplified model specified as active bins ~ extension + (1|block:replicate), with degrees of freedom obtained with the Kenward Roger approximation, and on sex-specific simplified models for mean peak activity hour, with model specified as mean peak hour ~ extension.

| **Response variable** | **Treatment (LED extension) comparison** | **Estimate (estimated marginal means, Tukey pairwise tests)** | **SE** | **Df** | **t-ratio** | **p-value** |
|---|---|---|---|---|---|---|
| Mean active 5-minute bins | BSW – no extension | 12.65 | 6.23 | 211 | 2.03 | 0.18 |
| | BSW – RB | 10.16 | 6.29 | 211 | 1.62 | 0.37 |
| | BSW – RGB | 18.36 | 6.24 | 211 | 2.94 | 0.019 |
| | No extension – RB | -2.49 | 5.88 | 210 | -0.42 | 0.97 |
| | No extension – RGB | 5.71 | 5.83 | 210 | 0.98 | 0.76 |
| | RB – RGB | 8.20 | 5.89 | 211 | 1.39 | 0.51 |
| **Response variable** | **Treatment (LED extension) comparison** | **Estimate (Tukey pairwise differences)** | **SE** | **t-value** | **p-value** | |
| Mean peak hour, females | No extension – BSW | -4.11 | 0.86 | -4.77 | <0.001 | |
| | RB – BSW | -1.37 | 0.84 | -1.63 | 0.37 | |
| | RGB – BSW | -0.34 | 0.86 | -0.40 | 0.98 | |
| | RB – no extension | 2.74 | 0.85 | 3.23 | 0.0086 | |
| | RGB – no extension | 3.77 | 0.86 | 4.37 | <0.001 | |
| | RGB-RB | 1.03 | 0.84 | 1.22 | 0.61 | |
| Mean peak hour, males | No extension – BSW | 0.078 | 1.40 | 0.056 | 1.00 | |
| | RB – BSW | 4.12 | 1.45 | 2.84 | 0.028 | |
| | RGB – BSW | 4.61 | 1.41 | 3.26 | 0.0081 | |
| | RB – no extension | 4.04 | 1.23 | 3.29 | 0.0073 | |
| | RGB – no extension | 4.53 | 1.18 | 3.84 | 0.0013 | |
| | RGB – RB | 0.49 | 1.24 | 0.39 | 0.98 | |
| **Response variable** | **Comparison** | **Dunn test statistic** | | **p-value** | | |
| Active lifespan (time until detected movement ceased) | No extension – BSW | -1.88 | | 0.12 | | |
| | RB – BSW | -3.51 | | 0.0027 | | |
| | RB – no extension | -1.63 | | 0.16 | | |
| | RGB – BSW | -3.42 | | 0.0019 | | |
| | RGB – no extension | -1.55 | | 0.15 | | |
| | RGB – RB | 0. 074 | | 0.94 | | |
